# Common ALS/FTD risk variants in *UNC13A* exacerbate its cryptic splicing and loss upon TDP-43 mislocalization

**DOI:** 10.1101/2021.04.02.438170

**Authors:** Anna-Leigh Brown, Oscar G. Wilkins, Matthew J. Keuss, Sarah E. Hill, Matteo Zanovello, Weaverly Colleen Lee, Flora C.Y. Lee, Laura Masino, Yue A. Qi, Sam Bryce-Smith, Alexander Bampton, Ariana Gatt, Hemali Phatnani, NYGC ALS Consortium, Giampietro Schiavo, Elizabeth M.C. Fisher, Towfique Raj, Maria Secrier, Tammaryn Lashley, Jernej Ule, Emanuele Buratti, Jack Humphrey, Michael E. Ward, Pietro Fratta

## Abstract

Variants within the *UNC13A* gene have long been known to increase risk of amyotrophic lateral sclerosis (ALS) and frontotemporal dementia (FTD), two related neurodegenerative diseases defined by mislocalization of the RNA-binding protein TDP-43. Here, we show that TDP-43 depletion induces robust inclusion of a cryptic exon (CE) within *UNC13A*, a critical synaptic gene, resulting in nonsense-mediated decay and protein loss. Strikingly, two common polymorphisms strongly associated with ALS/FTD risk directly alter TDP-43 binding within the CE or downstream intron, increasing CE inclusion in cultured cells and in patient brains. Our findings, which are the first to demonstrate a genetic link specifically between loss of TDP-43 nuclear function and disease, reveal both the mechanism by which *UNC13A* variants exacerbate the effects of decreased nuclear TDP-43 function, and provide a promising therapeutic target for TDP-43 proteinopathies.

**One-Sentence Summary:** Shared ALS/FTD risk variants increase the sensitivity of a cryptic exon in the synaptic gene *UNC13A* to TDP-43 depletion.

## Main Text

### Introduction

Amyotrophic lateral sclerosis (ALS) and frontotemporal dementia (FTD) are devastating adult-onset neurodegenerative disorders with shared genetic causes and common pathological aggregates (*1*–*3*). Genome-wide association studies (GWASs) have repeatedly demonstrated a shared risk locus between ALS and FTD within the crucial synaptic gene *UNC13A,* although the mechanism underlying this association has remained elusive (*4*).

ALS and FTD are pathologically defined by cytoplasmic aggregation and nuclear depletion of TAR DNA-binding protein 43 (TDP-43) in the vast majority (>97%) of ALS cases and in 45% of FTD cases (FTLD-TDP) (*5*). TDP-43, an RNA-binding protein (RBP), primarily resides in the nucleus and plays key regulatory roles in RNA metabolism, including acting as a splicing repressor. Upon TDP-43 nuclear depletion – an early pathological feature in ALS/FTLD-TDP – non-conserved intronic sequences are de-repressed and erroneously included in mature RNAs. These events are referred to as cryptic exons (CEs) and can lead to premature stop-codons/polyadenylation and transcript degradation (*6*, *7*). Recently, TDP-43 loss was found to induce a CE in the *Stathmin 2* (*STMN2*) transcript, which can serve as a functional readout for TDP-43 proteinopathy, as it appears selectively in affected patient tissue and its level correlates with TDP-43 phosphorylation (*8–10*).

In this study, we report a novel CE in *UNC13A* which promotes nonsense-mediated decay, and is present at remarkably high levels in patient neurons. Strikingly, we find that ALS/FTD risk-associated SNPs within *UNC13A* promote increased inclusion of this CE. We thus elucidate the molecular mechanism behind one of the top GWAS hits for ALS/FTD, and provide a promising new therapeutic target for TDP-43 proteinopathies.

## Results

### TDP-43 knockdown leads to inclusion of a cryptic exon in UNC13A

To discover novel CEs induced by TDP-43 depletion, we performed RNA-seq on human induced pluripotent stem cell (iPSC)-derived cortical-like i^3^Neurons in which we reduced TDP-43 expression through CRISPR inhibition (CRISPRi) (*10*–*13*). We identified 179 CEs, including several previously reported, such as *AGRN*, *PFKP* and *STMN2* (*6*–*9*) (**Fig. 1A**; data S1) (**Fig. 1B**; data S2). Interestingly, we observed robust mis-splicing in two members of the UNC13 synaptic protein family, *UNC13A* and *UNC13B* (**Fig. 1C-F**). Notably, *UNC13A* polymorphisms modify both disease risk and progression in ALS and FTLD-TDP (*4*, *14*–*21*) pointing towards a potential functional relationship between TDP-43, *UNC13A*, and disease risk.

**Fig. 1.**
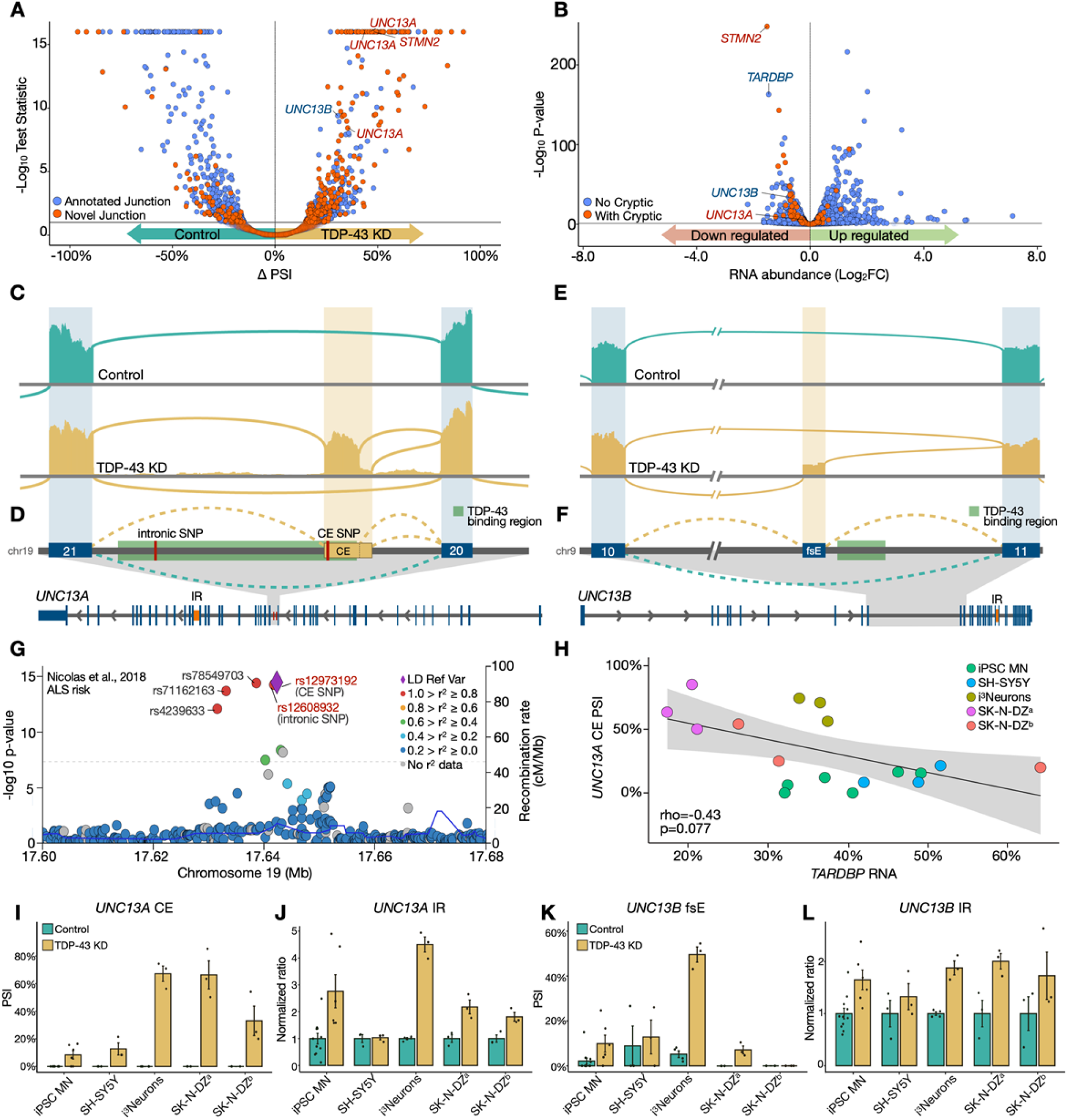
TDP-43 depletion in neurons leads to altered splicing in synaptic genes *UNC13A* and *UNC13B*. **(A)** Differential splicing and **(B)** expression in control (N=4) and CRISPRi TDP-43 depleted (N=3) iPSC-derived cortical-like i^3^Neurons. Each point denotes a splice junction (A) or gene (B). **(C)** Sashimi plots showing cryptic exon (CE) inclusion between exons 20 and 21 of *UNC13A* upon TDP-43 knockdown (KD). **(D,F)** Schematics showing intron retention (IR, lower schematic, orange), TDP-43 binding region (*22*)(green), and two ALS/FTLD associated SNPs (red). (**E)** Sashimi plot of *UNC13B* showing inclusion of the frameshifting exon (fsE) upon TDP-43 KD. (**G)** LocusZoom plot of the *UNC13A* locus in the latest ALS GWAS. Lead SNP *rs12973192* plotted as purple diamond, other SNPs coloured by linkage disequilibrium with *rs12973192* in European individuals from 1000 Genomes. **(H)** Correlation between relative *TARDBP* RNA and *UNC13A* CE PSI across five TDP-43 knockdown datasets **(I,K)** PSI of TDP-43 regulated splicing in *UNC13A* and *UNC13B* across neuronal datasets. **(J,L)** Intron retention ratio of TDP-43 regulated retained introns in *UNC13A* and *UNC13B* across neuronal datasets.

Inspection of the *UNC13A* gene revealed a previously unreported CE after TDP-43 knockdown (KD), with both a shorter and longer form, between exons 20 and 21 (**Fig. 1C**), and increased IR between exons 31 and 32 (fig. S1B). One ALS/FTLD-TDP risk SNP – *rs12973192* (*17*) – lies 16 bp inside the CE (henceforth referred to as the CE SNP). Another SNP – *rs12608932* (*4*) *–* is located 534 bp downstream of the donor splice site of the CE inside the same intron (henceforth referred to as the intronic SNP) (**Fig. 1D**). While there are five polymorphisms associated with ALS risk along *UNC13A*, they are all in high linkage disequilibrium (LD) in European populations with both the CE and intronic SNPs, and are present in 35% of individuals (**Fig. 1G**)(*17*). The close proximity of the disease-associated SNPs to the *UNC13A* CE suggests that the SNPs may influence *UNC13A* splicing. In *UNC13B*, TDP-43 KD led to the inclusion of an annotated frame-shift-inducing exon between exons 10 and 11, henceforth referred to as the *UNC13B* frameshift exon (fsE), and increased intron retention (IR) between exon 21 and 22 (**Fig. 1E,F**; fig. S1A).

In support of a direct role for TDP-43 regulation of *UNC13A* and *UNC13B*, we found multiple TDP-43 binding peaks both downstream and within the body of the *UNC13A* CE (**Fig. 1D**) and IR (fig. S1B) (*22*), and *UNC13A* CE inclusion negatively correlated with *TARDBP* RNA levels (rho = −0.43, p=0.077, **Fig. 1H**). Additionally, TDP-43 binding peaks were present near both splice events in *UNC13B* (**Fig. 1F**; fig. S1A) (*22*). We also detected these splicing changes in RNA-seq from TDP-43 depleted SH-SY5Y and SK-N-DZ neuronal lines, as well as in publicly available iPSC-derived motor neurons (MNs) (*8*) and SK-N-DZ datasets (*23*)(**Fig. 1I-L**; fig. S1C), and validated them by PCR in SH-SY5Y and SK-N-DZ cell lines (fig. S1D,E).

### UNC13A and UNC13B RNA and protein are downregulated by TDP-43 knockdown

Next, we examined whether incorrect splicing of *UNC13A* and *UNC13B* affected transcript levels in neurons and neuron-like cells. TDP-43 KD significantly reduced *UNC13A* RNA abundance in the three cell types with the highest levels of cryptic splicing (FDR < 0.1; **Fig. 2A**, **Fig. 1I**). Likewise, *UNC13B* RNA was significantly downregulated in four datasets (FDR < 0.1) (**Fig. 2B**). We confirmed these results by qPCR in SH-SY5Y and SK-N-DZ cell lines (fig. S2A). The number of ribosome footprints aligning to *UNC13A* and *UNC13B* was reduced after TDP-43 KD (**Fig. 2C**; fig. S2B, data S3). TDP-43 KD also decreased expression of UNC13A and UNC13B at the protein level, as assessed by quantitative proteomics with liquid chromatography tandem-mass spectrometry and western blot (**Fig. 2D,E**). These data suggest that the mis-splicing in *UNC13A* and *UNC13B* after TDP-43 KD reduces their transcript and protein abundance in neurons.

**Fig. 2.**
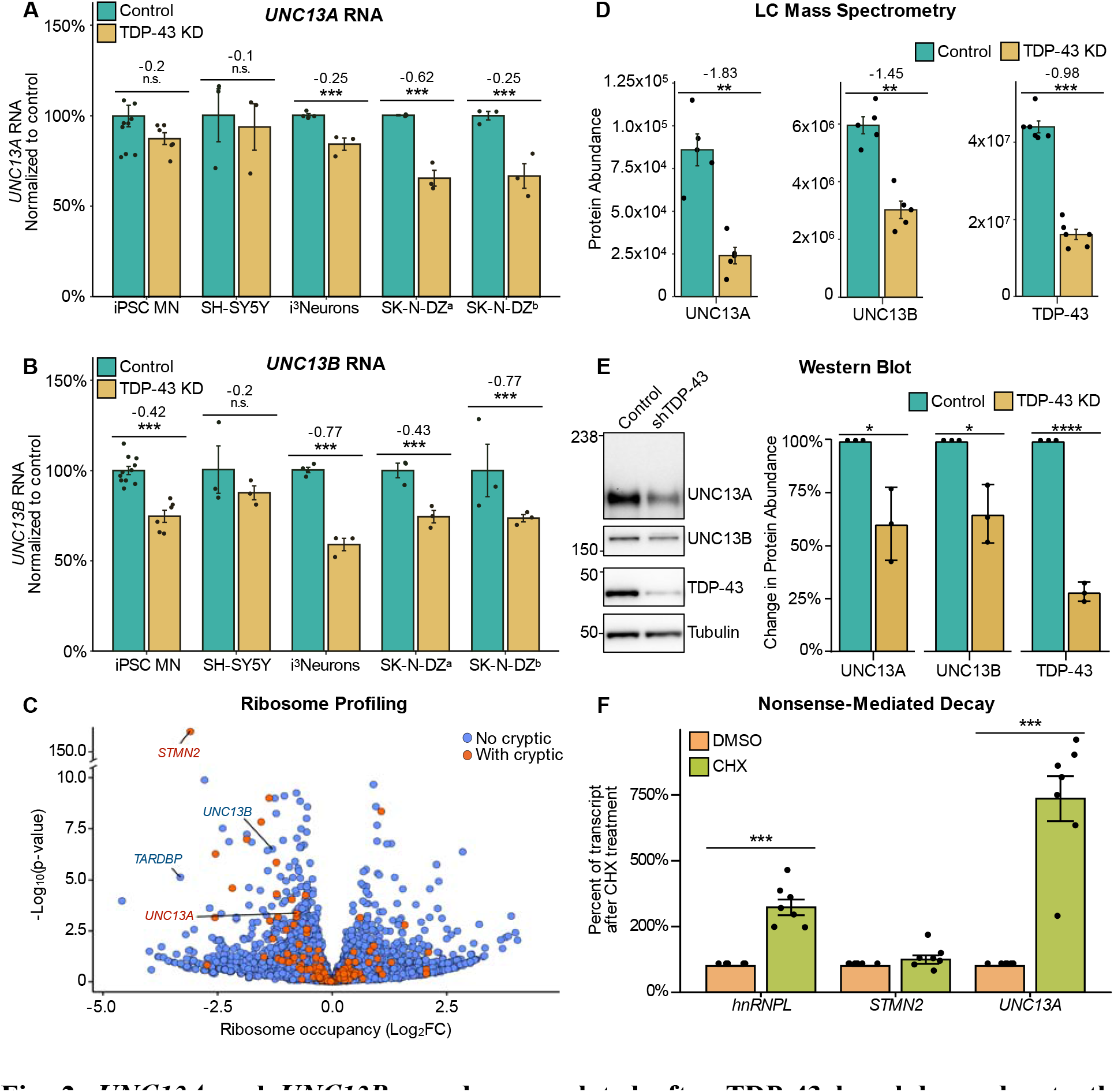
*UNC13A* and *UNC13B* are downregulated after TDP-43 knockdown due to the production of NMD-sensitive transcripts. Relative gene expression levels for *UNC13A **(A)*** and *UNC13B* **(B)** after TDP-43 knockdown across neuronal cell lines. Normalized RNA counts are shown as relative to control mean. Numbers show log_2_ fold change calculated by DESeq2. Significance shown as adjusted p-values from DESeq2. **(C)** Ribosome profiling of TDP-43 knockdown in i^3^Neurons shows reduction in ribosome occupancy of *STMN2*, *UNC13A* and *UNC13B* transcripts. **(D)** Mass spectrometry-based proteomic analysis shows reduction in protein abundance of UNC13A, UNC13B and TDP-43 upon TDP-43 knockdown in i^3^Neurons. Numbers refer to log_2_ fold change of unique peptide fragments, P-values from Wilcoxon test. **(E)** Western blot analysis of protein lysates from untreated and TDP-43 knockdown SH-SY5Y cells show a significant reduction in UNC13A and UNC13B proteins levels after TDP-43 depletion. Graphs represent the means ± S.E., N=3, One sample t-test, **(F)** Transcript expression upon CHX treatment suggests *UNC13A* but not *STMN2*, are sensitive to nonsense-mediated decay. *HNRNPL* (heterogeneous nuclear ribonucleoprotein L) is a positive control. Significance levels reported as * (p<0.05) ** (p<0.01) *** (p<0.001) **** (p <0.0001).

The *UNC13A* CE contains a premature termination codon (PTC) and is thus predicted to promote nonsense-mediated decay (NMD). Cycloheximide (CHX) treatment, which stalls translation and impairs NMD, increased CE inclusion in *UNC13A* after TDP-43 KD. Conversely, CHX did not alter levels of the aberrant *STMN2* transcript, which is not predicted to undergo NMD (**Fig. 2F**). Taken together, our data suggests that TDP-43 is critical for maintaining normal expression of the presynaptic proteins *UNC13A* and *UNC13B* by ensuring their correct pre-mRNA splicing.

### UNC13A cryptic exon is highly expressed in TDP-43-depleted patient neurons

To explore whether the *UNC13A* CE could be detected in patient tissues affected by TDP-43 pathology, we first analysed RNA-seq from neuronal nuclei sorted from frontal cortices of ALS/FTLD patients (*24*). We compared levels of *UNC13A* CE to levels of a CE in *STMN2* known to be regulated by TDP-43. Both *STMN2* and *UNC13A* CEs were exclusive to TDP-43-depleted nuclei, and, strikingly, in some cases the *UNC13A* CE percent spliced in (PSI) reached 100% (**Fig. 3A**). This suggests that in patients there will be a significant loss of UNC13A expression within the subpopulation of neurons with TDP-43 pathology.

**Fig. 3.**
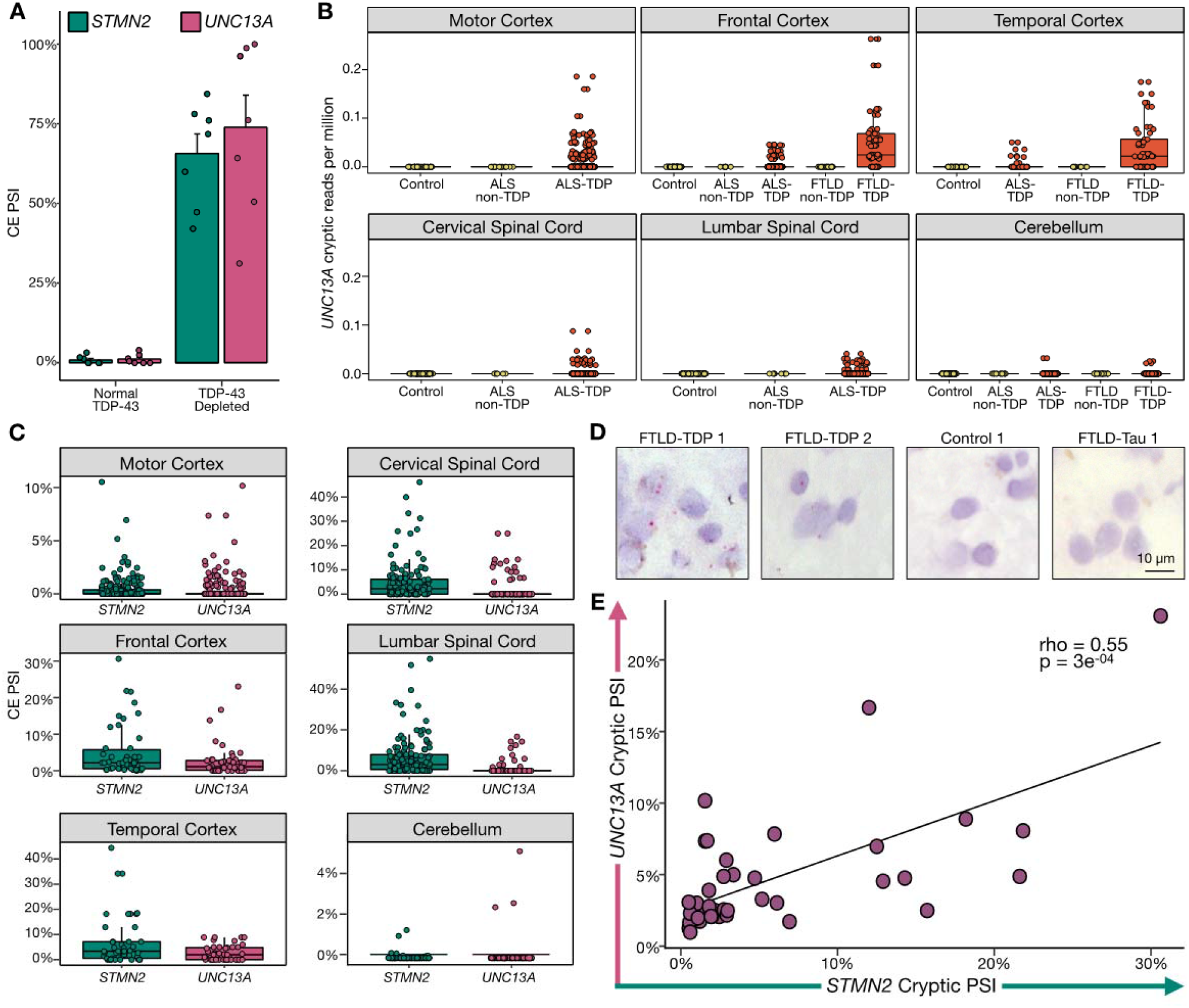
*UNC13A* CE is highly expressed in ALS/FTLD patient tissue and correlates with known markers of TDP-43 loss of function. **(A)***UNC13A* and *STMN2* CE expression in ALS/FTLD patient frontal cortex neuronal nuclei from (*24*) sorted according to the expression of nuclear TDP-43. **(B)***UNC13A* CE expression in bulk RNA-seq from NYGC ALS Consortium normalized by library size across disease and tissue samples. ALS cases stratified by mutation status, FTLD cases stratified by pathological subtype. **(C)** CE expression throughout ALS/FTLD-TDP cases across tissue **(D)** BaseScope detection of *UNC13A* CE (red foci) in FTLD-TDP but not control or FTLD-Tau frontal cortex samples. **(E)** Correlation in ALS/FTLD-TDP cortex between *UNC13A* and *STMN2* CE PSI in patients with at least 30 spliced reads across the CE locus.

Next, we quantified *UNC13A* CE inclusion in bulk RNA-seq from the NYGC ALS Consortium, a dataset containing 1,349 brain and spinal cord tissues from a total of 377 ALS, FTLD, and control individuals. The *UNC13A* CE was detected exclusively in FTLD-TDP and ALS-TDP cases (89% and 38% respectively), with no detection in ALS-non-TDP (*SOD1* and *FUS* mutations), FTLD-non-TDP (FTLD-TAU and FTLD-FUS), or control cases. The lower detection rate in ALS versus FTLD is likely due to the lower expression of *UNC13A* in the spinal cord (fig. S3A). Thus, pathological *UNC13A* CEs occur *in vivo* and are specific to neurodegenerative disease subtypes in which mislocalization and nuclear depletion of TDP-43 occurs.

*UNC13A* CE expression mirrored the known tissue distribution of TDP-43 aggregation and nuclear clearance (*25*): it was specific to ALS-TDP spinal cord and motor cortex, as well as FTLD-TDP frontal and temporal cortices, but absent from the cerebellum in disease and control states (**Fig. 3B**). Despite the CE PSI being diluted by both the presence of unaffected cells and NMD in bulk RNA-seq, we were still able to detect CE above 20% in some samples. Furthermore, although, unlike the STMN2 CE, the *UNC13A* CE induces NMD, it was detected at similar levels to *STMN2* CE in cortical regions, whilst *STMN2* CE was more abundant in the spinal cord (**Fig. 3C**). We next investigated whether *UNC13A* CEs could be visualised by *in situ* hybridisation (ISH) in FTLD patient brains. Using a probe targeting the *UNC13A* CE on frozen frontal cortex tissue, we detected staining significantly above background in 4 out of 5 tested FTLD-TDP cases, but in none of the FTLD-Tau (n=3) or control (n=5) cases (**Fig. 3D**; fig. S3B). To assess if *UNC13A* CE levels in bulk tissue was related to the level of TDP-43 proteinopathy, we used *STMN2* CE PSI as a proxy, as it correlates with the burden of phosphorylated TDP-43 in patient samples (*10*). As expected, across the NYGC ALS Consortium samples we observed a significant positive correlation between the level of *STMN2* CE PSI and *UNC13A* CE PSI (rho = 0.55, p = 3.0e-4) (**Fig. 3E**). Combined, our analysis reveals a strong relationship between TDP-43 pathology and *UNC13A* CE levels, supporting a model for direct regulation of *UNC13A* mRNA splicing by TDP-43 in patients.

### rs12973192(G) and rs12608932(C) combine to promote cryptic splicing

To test whether the ALS/FTD *UNC13A* risk SNPs promote cryptic splicing, which could explain their link to disease, we assessed *UNC13A* CE levels across different genotypes, and found significantly increased levels in cases homozygous for CE *rs12973192(G)* and intronic *rs12608932(C)* SNPs (fig. S4A-B). To ensure that this was not simply due to more severe TDP-43 pathology in these samples, we normalised by the level of *STMN2* cryptic splicing, and again found a significantly increased level of the *UNC13A* CE in cases with homozygous risk variants (Wilcoxon test, p < 0.001) (**Fig. 4A**; fig. S4C,D). Next, we performed targeted RNA-seq on *UNC13A* CE from temporal cortices of ten heterozygous risk allele cases and four controls. We detected significant biases towards reads containing the risk allele (p < 0.05, single-tailed binomial test) in six samples, with a seventh sample approaching significance (**Fig. 4B**), suggesting that the two ALS/FTLD-linked variants promote cryptic splicing *in vivo*.

**Fig. 4.**
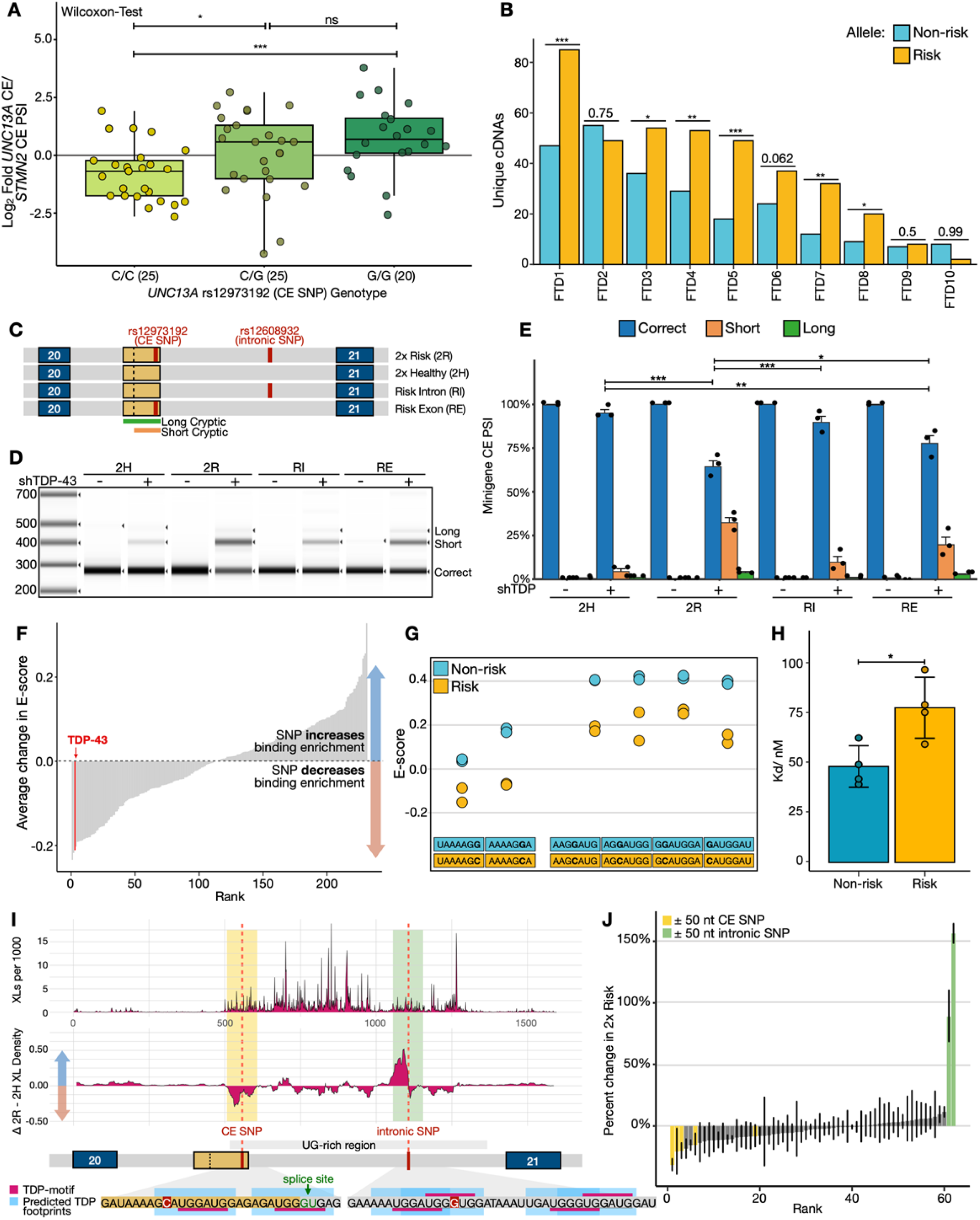
*UNC13A* ALS/FTD risk variants enhance *UNC13A* CE splicing in patients and in vitro by altering TDP-43 pre-mRNA binding. **(A)** Ratio *UNC13A / STMN2* CE PSI, split by genotype for *UNC13A* risk alleles. **(B)** Unique cDNAs from targeted RNA-seq in ten CE SNP heterozygous FTLD-TDP patients. p-values from single-tailed binomial tests. FTD1, 5, and 7 are *C9orf72 hexanucleotide repeat* carriers **(C)** Illustration of *UNC13A* minigenes containing exon 20, intron 20, and exon 21 with both risk SNPs (2R), both healthy SNPs (2H), or risk SNP in CE (RE) or intron (RI). **(D)** Representative image of RT-PCR products from *UNC13A* minigenes in SH-SY5Y ± TDP-43 KD. **(E)** Quantification of (D) plotted as means ± S.E. N=3, One-way ANOVA analysis; **(F)** Average change in E-value (measure of binding enrichment) across proteins for heptamers containing risk/healthy CE SNP allele; red - TDP-43. **(G)** Each CE SNP heptamer’s TDP-43 E-value. **(H)** Binding affinities between TDP-43 and 14-nt RNA containing the healthy or risk sequence measured by ITC; 4 replicates. **(I)** TDP-43 iCLIP of 2R and 2H minigenes: top - average crosslink density; bottom - average density change 2R - 2H (rolling window = 20 nt, units = crosslinks per 1,000). Cartoon - predicted TDP-43 binding footprints (UGNNUG motif). **(J)** Fractional changes at iCLIP peaks for 2R versus 2H minigene (mean and 75% confidence interval shown). Peaks that are within 50nt of each SNP are highlighted. *** (p<0.001) ** (p<0.01) * (p<0.05).

To specifically examine whether the CE or the intronic SNP of *UNC13A* promote CE splicing, we generated four variants of minigenes containing *UNC13A* exon 20, intron 20, and exon 21, featuring both risk alleles (2R), both non-risk alleles (2H), the risk allele within the CE (*rs12973192*) (RE), or the risk allele in the intron (*rs12608932*) (RI) (**Fig. 4C**). We then expressed these minigenes in SH-SY5Y cells with doxycycline-inducible TDP-43 knockdown. We found that both the CE SNP and, to a lesser extent, intronic SNP independently promoted CE inclusion, with the greatest overall levels detected for the 2R minigene (**Fig. 4D,E**).

To explore how these two SNPs might act to enhance CE splicing, we analyzed a dataset of *in vitro* RNA heptamer/RBP binding enrichments, and examined the effect of the SNPs on relative RBP enrichment (*26*). Strikingly, when investigating which RBPs were most impacted in their RNA binding enrichment by the CE-risk SNP, TDP-43 had the third largest decrease of any RBP, with only two non-human RBPs showing a larger decrease (**Fig. 4F,G**; fig. S4E,F). To test whether the CE SNP directly inhibited *in vitro* TDP-43 binding, we performed isothermal titration calorimetry using recombinant TDP-43 and 14-nt RNAs. As predicted, we observed an increased Kd for RNA containing the CE risk SNP (**Fig. 4H;** fig. S4G,H; Data S4). Together these data predict that the *UNC13A* CE SNP may directly inhibit TDP-43 binding.

To directly study the impact of the SNPs on TDP-43 binding to *UNC13A* pre-mRNA, we performed TDP-43 iCLIP with cells expressing either the 2R or 2H minigene. We observed a striking enrichment of crosslinks within the ~800nt UG-rich region containing both SNPs in intron 20 (**Fig. 4I**). When comparing the 2R with the 2H minigene, the peaks with the largest fractional changes were in close proximity of each SNP; similarly, we detected a 21% decrease in total TDP-43 crosslinks centred around the CE SNP and a 73% increase upstream of the intronic SNP (**Fig. 4I, J**, fig. S4I; 50 nucleotide windows). These data demonstrate that these two disease-risk SNPs distort the pattern of TDP-43/RNA interactions, decreasing TDP-43 binding near the CE donor splice site, thus exacerbating *UNC13A* CE inclusion upon nuclear TDP-43 depletion.

## Discussion

Our results support a model wherein TDP-43 nuclear depletion and the intronic and CE SNPs in *UNC13A* synergistically reduce expression of *UNC13A*, a gene that is critical for normal neuronal function. In this model, when nuclear TDP-43 levels are normal in healthy individuals, TDP-43 efficiently binds to *UNC13A* pre-mRNA and prevents CE splicing, regardless of *UNC13A* SNPs. Conversely, severe nuclear depletion of TDP-43 in end-stage disease induces CE inclusion in all cases. However, the common intronic and CE SNPs in *UNC13A* alter TDP-43 binding to *UNC13A* pre-mRNA and may make *UNC13A* CE more sensitive to partial TDP-43 loss that occurs early in degenerating neurons, explaining their associated risk effect. Strikingly, we found that both risk alleles for these SNPs independently and additively promoted cryptic splicing *in vitro.* Intriguingly, when the two variants are not co-inherited, as seen in East Asian individuals with ALS, an attenuated effect is observed (*20*). A similar phenomenon wherein SNP pairs both contribute to risk has been widely studied at the APOE locus in Alzheimer’s disease (*27*).

UNC13-family proteins are highly conserved across metazoans and are essential for calcium-triggered synaptic vesicle release (*28*). In mice, double knockout of *UNC13A* and *UNC13B* inhibits both excitatory and inhibitory synaptic transmission in hippocampal neurons and greatly impairs transmission at neuromuscular junctions (*29*, *30*). In TDP-43-negative neuronal nuclei derived from patients, the *UNC13A* CE is present in up to 100% of transcripts, suggesting that expression of functional *UNC13A* is greatly reduced, which could impact normal synaptic transmission.

TDP-43 loss induces hundreds of splicing changes, a number of which have also been detected in patient brains. However, it has remained unclear whether these events – even those that occur in crucial neuronal genes – contribute to disease pathogenesis. That genetic variation influencing the *UNC13A* CE inclusion can lead to changes in ALS/FTD susceptibility and progression strongly supports *UNC13A* downregulation to be one of the critical consequences of TDP-43 loss of function. Excitingly, *UNC13A* provides a generalizable therapeutic target for 97% of ALS and approximately half of FTD cases. These findings are also of interest to other neurodegenerative diseases, such as Alzheimer’s disease, Parkinson’s disease and chronic traumatic encephalopathy, in which TDP-43 depletion is also observed in a significant fraction of cases.

## Acknowledgments

We thank Frédéric Allain for the His-tagged TDP-43 plasmid, Cristiana Stuani for guidance on TDP-43 purification, and Martina Hallegger for guidance on TDP-43 iCLIP.

## Funding

UK Medical Research Council [MR/M008606/1 and MR/S006508/1] (PF)

UK Motor Neurone Disease Association (PF)

Rosetrees Trust (PF,AG)

UCLH NIHR Biomedical Research Centre (PF) 4 Year Wellcome Trust Studentship (OGW)

European Union’s Horizon 2020 research and innovation programme (835300-RNPdynamics) (JU)

Cancer Research UK (FC001002) (JU)

UK Medical Research Council (FC001002) (JU) Wellcome Trust (FC001002) (JU)

Collaborative Center for X-linked Dystonia-Parkinsonism (WCL, EMCF)

Intramural Research Program of the National Institutes of Neurological Disorders and Stroke, NIH, Bethesda, MD (MEW,SEH)

Chan Zuckerberg Initiative (MEW)

The Robert Packard Center for ALS Research (MEW)

E.B. is funded by AriSLA PathensTDP project (EB)

Wolfson Foundation (AB)

Brightfocus Foundation postdoctoral research fellowship (SEH)

Wellcome Trust Investigator Award (107116/Z/15/Z) (GS)

UK Dementia Research Institute Foundation award (UKDRI-1005)

(GS) Alzheimer’s Research UK senior fellowship (TL)

Alzheimers Society (AG)

UKRI Future Leaders Fellowship (MR/T042184/1) (MS)

## Author contributions

Conceptualization: ALB,OGW,MJK,SEH,JH,MEW,PF

Data curation: ALB,OGW,MZ,SBS

Formal analysis: ALB,OGW,MJK,MZ,SBS,AB

Funding acquisition: PF,MEW,EB

Investigation: ALB,OGW,MJK,SEH,MZ,FCYL,LM,YAQ,SBS,AB,WCL,AG

Methodology: ALB,OGW,MJK,SEH,JH,MEW,PF

Project administration: PF,MEW

Resources: HP,TL,EB

Software: ALB,OGW,MZ,SBS,JH

Supervision: PF,MEW,JH,JU,MS,TR,TL,EMCF,GS

Visualization: ALB,OGW,MJK,WCL

Writing – original draft: ALB,OGW,MJK,MEW,PF

Writing – review & editing: SEH,WCL,EB,JU,JH

## Data and materials availability

Analysis code and data to reproduce figures available: https://github.com/frattalab/unc13a_cryptic_splicing/

RNA-Seq Data for i3Neurons, SH-SY5Y and SK-N-DZ^a^ are available through the European Nucleotide Archive (ENA) under accession PRJEB42763.

Public data was obtained from Gene Expression Omnibus (GEO): iPSC MNs (Klim et al., 2019)-GSE121569, SK-N-DZ^b^-GSE97262, and FACS-sorted frontal cortex neuronal nuclei-GSE126543.

*Riboseq:* E-MTAB-10235.

*Targeted RNA seq:* E-MTAB-10237

*Minigene iCLIP*: E-MTAB-10297

*NYGC ALS Consortium RNA-seq:* RNA-Seq data generated through the NYGC ALS Consortium in this study can be accessed via the NCBI’s GEO database (GEO GSE137810, GSE124439, GSE116622, and GSE153960). All RNA-Seq data generated by the NYGC ALS Consortium are made immediately available to all members of the Consortium and with other consortia with whom we have a reciprocal sharing arrangement. To request immediate access to new and ongoing data generated by the NYGC ALS Consortium and for samples provided through the Target ALS Postmortem Core, complete a genetic data request form at.

*NYGC ALS Consortium Whole Genome Seq:* to be released later with companion manuscript.

## Materials and Methods

### Human iPSC culture

All policies of the NIH Intramural research program were followed for the procurement and use of induced pluripotent stem cells (iPSCs). The iPSCs used in this study were from the WTC11 line, derived from a healthy thirty-year old male, and obtained from the Coriell cell repository. All culture procedures were conducted as previously (*11*). In short, iPSCs were grown on tissue culture dishes coated with hESC-qualified matrigel (Corning, REF 354277). They were maintained in Essential 8 Medium (E8; Thermo Fisher Scientific, Cat. No. A1517001) supplemented with 10 μM ROCK inhibitor (RI; Y-27632; Selleckchem, Cat. No. S1049) in a 37°C, 5% CO_2_ incubator. Media was replaced every 1-2 days as needed. Cells were passaged with accutase (Life Technologies, Cat. No. A1110501), 5-10 minutes treatment at 37°C. Accutase was removed and cells were washed with PBS before re-plating. Following dissociation, cells were plated in E8 media supplemented with 10 μM RI to promote survival. RI was removed once cells grew into colonies of 5-10 cells.

The following cell lines/DNA samples were obtained from the NIGMS Human Genetic Cell Repository at the Coriell Institute for Medical Research: GM25256

### TDP-43 knockdown in human iPSCs

The human iPSCs used in this study were previously engineered (*11*, *13*) to express mouse Neurogenin-2 (NGN2) under a doxycycline-inducible promoter integrated at the AAVS1 safe harbor, as well as an enzymatically dead Cas9 (+/− CAG-dCas9-BFP-KRAB) integrated at a safe harbor at the CLYBL promoter (*12*).

To achieve knockdown, sgRNAs targeting either *TARDBP*/TDP-43 or a non-targeting control guide were delivered to iPSCs by lenti-viral transduction. To make the virus, Lenti-X Human Embryonic Kidney (HEK) cells were transfected with the sgRNA plasmids using Lipofectamine 3000 (Life Technologies, Cat. No. L3000150), then cultured for 2-3 days in the following media: DMEM, high glucose GlutaMAX Supplement media (Life Technologies, Cat. No. 10566024) with 10% FBS (Sigma, Cat. No. TMS-013-B), supplemented with viral boost reagent (ALSTEM, Cat. No. VB100). Virus was then concentrated from the media 1:10 in PBS using Lenti-X concentrator (Takara Bio, Cat. No. 631231), aliquoted and stored at −80°C for future use.

The sgRNAs were cloned into either pU6-sgRNA EF1Alpha-puro-T2A-BFP vector (gift from Jonathan Weissman; Addgene #60955) (*12*, *31*) or a modified version containing a human U6 promoter, a blasticidin (Bsd) resistance gene, and eGFP. sgRNA sequences were as follows: non-targeting control: GTCCACCCTTATCTAGGCTA and *TARDBP*: GGGAAGTCAGCCGTGAGACC.

Virus was delivered to iPSCs in suspension following an accutase split. Cells were plated and cultured overnight. The following morning, cells were washed with PBS and media was changed to E8 or E8+RI depending on cell density. Two days post lentiviral delivery, cells were selected overnight with either puromycin (10 μg/ml) or blasticidin (100 μg/ml). iPSCs were then expanded 1-2 days before initiating neuronal differentiation. Knockdown efficiency was tested at iPSC and neuronal stages using immunofluorescence, QT-PCR and observed in RNA-seq data.

### iPSC-derived i^3^Neuron differentiation and culture

To initiate neuronal differentiation, 20-25 million iPSCs per 15 cm plate were individualized using accutase on day 0 and re-plated onto matrigel-coated tissue culture dishes in N2 differentiation media containing: knockout DMEM/F12 media (Life Technologies Corporation, Cat. No. 12660012) with N2 supplement (Life Technologies Corporation, Cat. No. 17502048), 1x GlutaMAX (Thermofisher Scientific, Cat. No. 35050061), 1x MEM nonessential amino acids (NEAA) (Thermofisher Scientific, Cat. No. 11140050), 10 μM ROCK inhibitor (Y-27632; Selleckchem, Cat. No. S1049) and 2 μg/mL doxycycline (Clontech, Cat. No. 631311). Media was changed daily during this stage.

On day 3 pre-neuron cells were replated onto dishes coated with freshly made poly-L-ornithine (PLO; 0.1 mg/ml; Sigma, Cat. No. P3655-10MG), either 96-well plates (50,000 per well), 6-well dishes (2 million per well), or 15 cm dishes (45 million per plate), in i^3^Neuron Culture Media: BrainPhys media (STEMCELL Technologies, Cat. No. 05790) supplemented with 1x B27 Plus Supplement (ThermoFisher Scientific, Cat. No. A3582801), 10 ng/mL BDNF (PeproTech, Cat. No. 450-02), 10 ng/mL NT-3 (PeproTech, Cat. No. 450-03), 1 μg/mL mouse laminin (Sigma, Cat. No. L2020-1MG), and 2 ug/mL doxycycline (Clontech, Cat. No. 631311). i^3^Neurons were then fed three times a week by half media changes. i^3^Neuron were then harvested on day 17 post addition of doxycycline or 14 days after re-plating.

### Generation of Stable TDP-43 knockdown cell line

SH-SY5Y and SK-N-DZ cells were transduced with SmartVector lentivirus (V3IHSHEG_6494503) containing a doxycycline-inducible shRNA cassette for TDP-43. Transduced cells were selected with puromycin (1 μg/mL) for one week.

### Depletion of TDP-43 from immortalised human cell lines

SH-SY5Y cells for RT-qPCR validations and western blots were grown in DMEM/F12 containing Glutamax (Thermo) supplemented with 10% FBS (Thermo). For induction of shRNA against TDP-43 cells were treated with 5 μg/mL Doxycyline Hyclate (Sigma D9891). After 3 days media was replaced with Neurobasal (Thermo) supplemented with B27 (Thermo) to induce differentiation. After a further 7 days, cells were harvested for protein or RNA. SH-SY5Y and SK-N-DZ cells for RNA-seq experiments were treated with siRNA, as previously described (*23*).

### RNA-sequencing, differential gene expression and splicing analysis

For RNA-seq experiments of i^3^Neurons, the i^3^Neurons were grown on 96-well dishes. To harvest on day 17, media was completely removed, and wells were treated with tri-reagent (100 μL per well) (Zymo research corporation, Cat. No. R2050-1-200). Then 5 wells were pooled together for each biological replicate: control (n=3); TDP-43 knockdown (n=4). To isolate RNA, we used a Direct-zol RNA miniprep kit (Zymo Research Corporation, Cat. No. R2052), following manufacturer’s instructions including the optional DNAse step. Note: one control replicate did not pass RNA quality controls and so was not submitted for sequencing. Total RNA was then enriched for polyA and sequenced 2×75 bp on a HiSeq 2500 machine.

Samples were quality trimmed using Fastp with the parameter “qualified_quality_phred: 10”, and aligned to the GRCh38 genome build using STAR (v2.7.0f) (*32*) with gene models from GENCODE v31 (*33*). Gene expression was quantified using FeatureCounts (*34*) using gene models from GENCODE v31. Any gene which did not have an expression of at least 0.5 counts per million (CPM) in more than 2 samples was removed. For differential gene expression analysis, all samples were run in the same manner using the standard DESeq2 (*35*) workflow without additional covariates, except for the Klim MNs dataset, where we included the day of differentiation. DESeq2’s median of ratios, which controls for both sequencing depth and RNA composition, was used to normalize gene counts. Differential expression was defined at a Benjamini-Hochberg false discovery rate < 0.1. Our alignment pipeline is implemented in Snakemake version 5.5.4 (*36*) and available at: https://github.com/frattalab/rna_seq_snakemake.

Differential splicing was performed using MAJIQ (v2.1) (*37*) using the GRCh38 reference genome. A threshold of 0.1 ΔPSI was used for calling the probability of significant change between groups. The results of the deltaPSI module were then parsed using custom R scripts to obtain a PSI and probability of change for each junction. Cryptic splicing was defined as junctions with PSI < 0.05 in control samples, ΔPSI > 0.1, and the junction was unannotated in GENCODE v31. Our splicing pipeline is implemented in Snakemake version 5.5.4 and available at: https://github.com/frattalab/splicing.

Counts for specific junctions were tallied by parsing the STAR splice junction output tables using bedtools (*38*). Splice junction parsing pipeline is implemented in Snakemake version 5.5.4 and available at: https://github.com/frattalab/bedops_parse_star_junctions

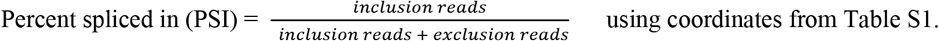

Intron retention was assessed using IRFinder (*39*) with gene models from GENCODE v31.

### Analysis of published iCLIP data

Cross-linked read files from TDP-43 iCLIP experiments in SH-SY5Y and human neuronal stem cells (*22*) were processed using iCount v2.0.1.dev implemented in Snakemake version 5.5.4, available at https://github.com/frattalab/pipeline_iclip. Sites of cross-linked reads from all replicates were merged into a single file using iCount group command. Significant positions of cross-link read density with respect to the same gene (GENCODE v34 annotations) were then identified using the iCount peaks command with default parameters. The pipeline

### Western Blot

SH-SY5Y cells were lysed directly in the sample loading buffer (Thermo NP0008). Lysates were heated at 95°C for 5 min with 100 mM DTT. If required lysates were passed through a QIAshredder (Qiagen) to shear DNA. Lysates were resolved on 4-12% Bis-Tris Gels (Thermo) or homemade 6% Bis-Tris gels and transferred to 0.45 μm PVDF (Millipore) membranes. After blocking with 5% milk, blots were probed with antibodies [Rb anti-UNC13A (Synaptic Systems 126 103); Rb anti-UNC13B (abcam ab97924); Rat anti-Tubulin (abcam ab6161), Mouse anti-TDP-43 (abcam ab104223)] for 2 hours at room temperature. After washing, blots were probed with HRP conjugated secondary antibodies and developed with Chemiluminescent substrate (Thermo) on a ChemiDoc Imaging System (Bio-Rad). Band intensity was measured with ImageJ (NIH).

### RT-qPCR

RNA was extracted from SH-SY5Y and SK-N-DZ cells with a RNeasy kit (Qiagen) using the manufacturer‘s protocol including the on column DNA digestion step. RNA concentrations were measured by Nanodrop and 1 μg of RNA was used for reverse transcription. First strand cDNA synthesis was performed with SSIV (Thermo 18090050) or RevertAid (Thermo K1622) using random hexamer primers and following the manufacturer’s protocol including all optional steps. Gene expression analysis was performed by qPCR using Taqman Multiplex Universal Master Mix (Thermo 4461882) and TaqMan assays (UNC13A-Fam Hs00392638_m1, UNC13B-Fam Hs01066405_m1, TDP-43-Vic Hs00606522_m1, GAPDH-Jun asay 4485713) on a QuantStudio 5 Real-Time PCR system (Applied Biosystems) and quantified using the ΔΔCt method (*40*).

### Nonsense-mediated decay (NMD) inhibition

Ten days post induction of shRNA against TDP-43 with 1 μg/ml doxycyline hyclate (Sigma D9891-1G), SH-SY5Y cells were treated either with 100 μM cycloheximide (CHX) or DMSO for 6 hours (*41*) before harvesting the RNA through RNeasy Minikit (Qiagen). Reverse transcription was performed using RevertAid cDNA synthesis kit (Thermo), and transcript levels were quantified by qPCR (QuantStudio 5 Real-Time PCR system, Applied Biosystems) using the ΔΔCt method and *GAPDH* as reference (*40*). Since it proved to undergo NMD (*42*), *hNRNPL* NMD transcript was used as a positive control.

### Quantification of TDP-43, UNC13A, and UNC13B using quantitative proteomics

i^*3*^Neurons were harvested from 6-well plates on day 17 post initiation of differentiation. Two wells were pooled for each biological replicate, n=6 for each control and TDP-43 knockdown neurons. To harvest, wells were washed with PBS, and then SP3 protein extraction was performed to extract intercellular proteins. Briefly, we harvested and lysed using a very stringent buffer (50 mM HEPES, 50 mM NaCl, 5 mM EDTA 1% SDS, 1% Triton X-100, 1% NP-40, 1% Tween 20, 1% deoxycholate and 1% glycerol) supplemental with cOmplete protease inhibitor cocktail at 1 tablet/10 ml ratio. The cell lysate was reduced by 10 mM dithiothreitol (30 min, 60°C) and alkylated using 20 mM iodoacetamide (30min, dark, room temperature). The denatured proteins were captured by hydrophilic magnetic beads, and tryptic on-beads digestion was conducted for 16 hours at 37°C. We injected 1 μg resulting peptides to a nano liquid chromatography (LC) for separation, and subsequently those tryptic peptides were analyzed on an Orbitrap Eclipse mass spectrometer (MS) coupled with a FAIMS interface using data-dependent acquisition (DDA) and data-independent acquisition (DIA). The peptides were separated on a 120 minute LC gradient with 2-35% solvent B (0.1% FA, 5% DSMO in acetonitrile), and FAIMS’s compensation voltages were set to −50, −65 and −80. For DDA, we used MS1 resolution at 12000 and cycle time was selected for 3 seconds, MS2 fragments were acquired by linear ion trap. For DIA, we used 8 m/z isolation windows (400-1000 m/z range), cycle time was set to 3 seconds, and MS2 resolution was set to 30000. The DDA and DIA MS raw files were searched against Uniprot-Human-Proteome_UP000005640 database with 1% FDR using Proteome Discoverer (v2.4) and Spectronaut (v14.1), respectively. The raw intensity of quantified peptides was normalized by total peptides intensity identified in the same sample. The DDA quantified TDP-43- and UNC13A-derived unique and sharing peptides were parsed out and used for protein quantification. Specifically, we visualized and quantified the unique peptides of UNC13A using their MS/MS fragment ion intensity acquired by DIA.

### Ribosome profiling

For ribosome profiling experiments, i^3^Neurons were grown on 15 cm plates, one plate per biological replicate for control (n=4) and TDP-43 knockdown (n=4) neurons. On day 17, i^3^Neuron Culture Medium was replaced 90 minutes prior to harvesting the neurons to boost translation. Then the medium was removed, cells were washed with cold PBS, PBS was removed and 900 μL of cold lysis buffer (20 mM Tris pH 7.4, 150 mM NaCl, 5mM MgCl_2_, 1 mM DTT freshly made, 100 ug/mL Cycloheximide, 1% TX100; 25 U/ml Turbo DNase I) was added to each 15 cm plate. Lysed cells were scraped and pipetted into microcentrifuge tubes on ice. Cells were then passed through a 26-gauge needle 10 times, and then centrifuged twice at 19,000xg 4°C, for 10 minutes, each time moving the supernatant to a fresh tube. Tubes containing supernatant were flash frozen in liquid nitrogen and stored at −80°C until processing.

Ribosome footprints from 3x TDP-43 knockdown and 3x control samples were generated and purified as described, using a sucrose cushion (McGlincy and Ingolia, 2017) and a customised library preparation method based on revised iCLIP (*43*). No rRNA depletion step was performed, and libraries were sequenced on an Illumina Hi-Seq 4000 machine (SR100). Reads were demultiplexed and adaptor/quality trimmed using Ultraplex (https://github.com/ulelab/ultraplex), then aligned with Bowtie2 against a reference file containing abundant ncRNAs that are common contaminants of ribosome profiling, including rRNAs (*44*). Reads that did not pre-map were then aligned against the human genome with STAR (*32*) and the resulting BAM files were deduplicated with UMI-tools (*45*). Multi-mapping reads were discarded and reads 28-30nt in length were selected for analysis. FeatureCounts (*34*) was used to count footprints aligning to annotated coding sequences, and DESEQ2 (*35*) was used for differential expression analysis, using default parameters in both cases. Periodicity analysis was performed using a custom R script, using transcriptome-aligned bam files. Raw data has been uploaded to E-MTAB-10235.

### Genome-wide association study data

Harmonised summary statistics for the latest ALS GWAS (*17*) were downloaded from the NHGRI-EBI GWAS Catalog (*46*) (accession GCST005647). Locus plots were created using LocusZoom (*47*), using linkage disequilibrium values from the 1000 Genomes European superpopulation (*48*).

### NYGC ALS Consortium RNA-seq cohort

Our analysis contains 377 patients with 1349 neurological tissue samples from the NYGC ALS dataset, including non-neurological disease controls, FTLD, ALS, FTD with ALS (ALS-FTLD), or ALS with suspected Alzheimer’s disease (ALS-AD). Patients with FTD were classified according to a pathologist’s diagnosis of FTD with TDP-43 inclusions (FTLD-TDP), or those with FUS or Tau aggregates. ALS samples were divided into the following subcategories using the available Consortium metadata: ALS with or without reported SOD1 or FUS mutations. All non-SOD1/FUS ALS samples were grouped as “ALS-TDP” in this work for simplicity, although reporting of postmortem TDP-43 inclusions was not systematic and therefore not integrated into the metadata. Confirmed TDP-43 pathology postmortem was reported for all FTLD-TDP samples.

Sample processing, library preparation, and RNA-seq quality control have been extensively described in previous papers (*10*, *49*). In brief, RNA was extracted from flash-frozen postmortem tissue using TRIzol (Thermo Fisher Scientific) chloroform, and RNA-Seq libraries were prepared from 500 ng total RNA using the KAPA Stranded RNA-Seq Kit with RiboErase (KAPA Biosystems) for rRNA depletion. Pooled libraries (average insert size: 375 bp) passing the quality criteria were sequenced either on an Illumina HiSeq 2500 (125 bp paired end) or an Illumina NovaSeq (100 bp paired end). The samples had a median sequencing depth of 42 million read pairs, with a range between 16 and 167 million read pairs.

Samples were uniformly processed, including adapter trimming with Trimmomatic and alignment to the hg38 genome build using STAR (2.7.2a) (*32*) with indexes from GENCODE v30. Extensive quality control was performed using SAMtools (*50*) and Picard Tools (*51*) to confirm sex and tissue of origin.

Uniquely mapped reads within the *UNC13A* locus were extracted from each sample using SAMtools. Any read marked as a PCR duplicate by Picard Tools was discarded. Splice junction reads were then extracted with RegTools (*52*) using a minimum of 8 bp as an anchor on each side of the junction and a maximum intron size of 500 kb. Junctions from each sample were then clustered together using LeafCutter (*53*) with relaxed junction filtering (minimum total reads per junction = 30, minimum fraction of total cluster reads = 0.0001). This produced a matrix of junction counts across all samples.

As the long CE acceptor was detected consistently in control cerebellum samples, as part of an unannotated cerebellum-enriched 35 bp exon containing a stop codon between exons 20 and 21 (sup fig 3C,D), we excluded the long CE acceptor for quantification of *UNC13A* CE PSI in patient tissue. Only samples with at least 30 spliced reads at the exon locus were included for correlations.

### BaseScope assay

Frozen tissue from the frontal cortex of FTLD-TDP (n = 5), FTLD-Tau (n = 3) and control (n = 3) cases were sectioned at 10 μm thickness onto Plus+Frost microslides (Solmedia). Immediately prior to use, sections were dried at RT and fixed for 15 minutes in pre-chilled 4 % paraformaldehyde. Sections were then dehydrated in increasing grades of ethanol and pre-treated with RNAscope^®^ hydrogen peroxide (10 mins, RT) and protease IV (30 mins, RT). The BaseScope™ v2-RED assay was performed using our UNC13A CE target probe (BA-Hs-UNC13A-O1-1zz-st) according to manufacturer guidelines with no modifications (Advanced Cell Diagnostics, Newark, CA). Sections were nuclei counterstained in Mayer’s haematoxylin (BDH) and mounted (VectaMount). Slides were also incubated with a positive control probe (Hs-PPIB-1 ZZ) targeting a common housekeeping gene and a negative control probe (DapB-1 ZZ) which targets a bacterial gene to assess background signal (< 1-2 foci per ~ 100 nuclei). Representative images were taken at x60 magnification.

Hybridised sections were graded, blinded to disease status, according to the relative frequency of red foci which should identify single transcripts with the *UNC13A* CE event. Grades were prescribed by relative comparison with the negative control slide. − = Less signal than negative control probe; + = similar signal strength to negative control; ++ = visibly greater signal than negative control, +++ = considerably greater signal than negative control.We identified a signal level above background (++ or +++) in 4 of 5 FTLD-TDP cases and a signal considerably above background (+++) level in 2 cases. All FTLD-Tau and control cases were graded as exhibiting either reduced (-) or comparable (+) signal relative to background.

### UNC13A genotypes in the NYGC ALS Consortium

Whole Genome Sequencing (WGS) was carried out for all donors, from DNA extracted from blood or brain tissue. Full details of sample preparation and quality control will be published in a future manuscript. Briefly, paired-end 150bp reads were aligned to the GRCh38 human reference using the Burrows-Wheeler Aligner (BWA-MEM v0.7.15) (*54*) and processed using the GATK best-practices workflow. This includes marking of duplicate reads by the use of Picard tools(*51*) (v2.4.1), followed by local realignment around indels, and base quality score recalibration using the Genome Analysis Toolkit (*55*, *56*) (v3.5). Genotypes for *rs12608932* and *rs12973192* were then extracted for the samples.

### Targeted RNA-seq

RNA was isolated from temporal cortex tissue of 10 FTLD-TDP and four control brains (6M, 4F, average age at death 70.6±5.8y, average disease duration 10.98±5.9y). 50 mg of flash-frozen tissue was homogenised in 700 μl of Qiazol (Qiagen) using a TissueRuptor II (Qiagen). Chloroform was added and RNA subsequently extracted following the spin-column protocol from the miRNeasy kit with DNase digestion (Qiagen). RNA was eluted off the column in 50 μl of RNAse-free water. RNA quantity and quality were evaluated using a spectrophotometer.

Purified RNA was reverse transcribed with Superscript IV (Thermo Fisher Scientific) using either sequence-specific primers containing sample-specific barcodes or random hexamers, following the manufacturer recommendations. Unique molecular identifiers (UMIs) and part of the P5 Illumina sequence were added either during first- or second-strand-synthesis (with Phusion HF 2x Master Mix) respectively. Barcoded primers were removed with exonuclease I treatment (NEB; 30 min) and subsequently bead/size selection of RT/PCR products (TotalPure NGS, Omega Biotek). Three rounds of nested PCR using Phusion HF 2x Master Mix (New England Biolabs) were used to obtain highly specific amplicons for the UNC13A cryptic, followed by gel extraction and a final round of PCR in which the full length P3/P5 Illumina sequences were added. Samples were sequenced with an Illumina HiSeq 4000 machine (SR100).

Raw reads were demultiplexed, adaptor/quality trimmed and UMIs were extracted with Ultraplex (https://github.com/ulelab/ultraplex), then aligned to the hg38 genome with STAR (*32*); for the hexamer data, a subsample of reads was used to reduce the number of PCR duplicates during analysis. Reads were deduplicated via analysis of UMIs with a custom R script; to avoid erroneous detection of UMIs due to sequencing errors, UMI sequences with significant similarity to greatly more abundant UMIs were discarded - this methodology was tested using simulated data, and final results were manually verified. Raw reads for targeted RNA-seq are available at E-MTAB-10237.

Primers used:

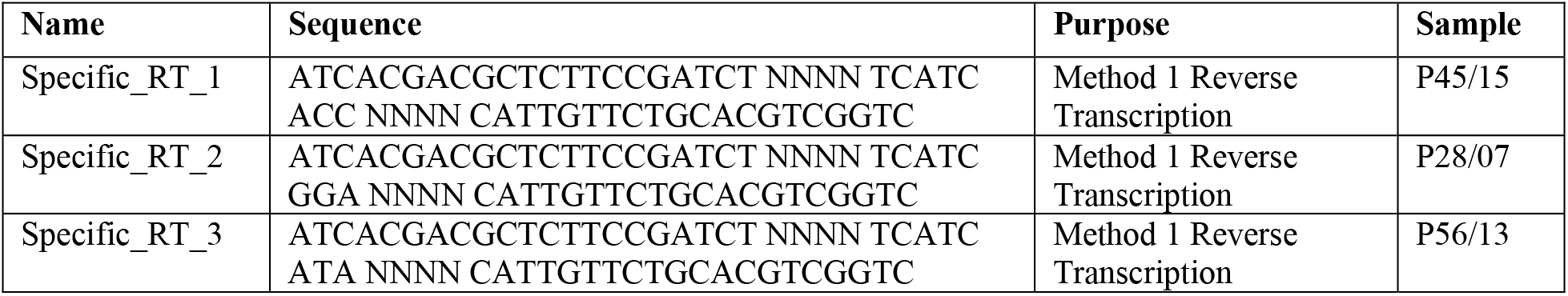
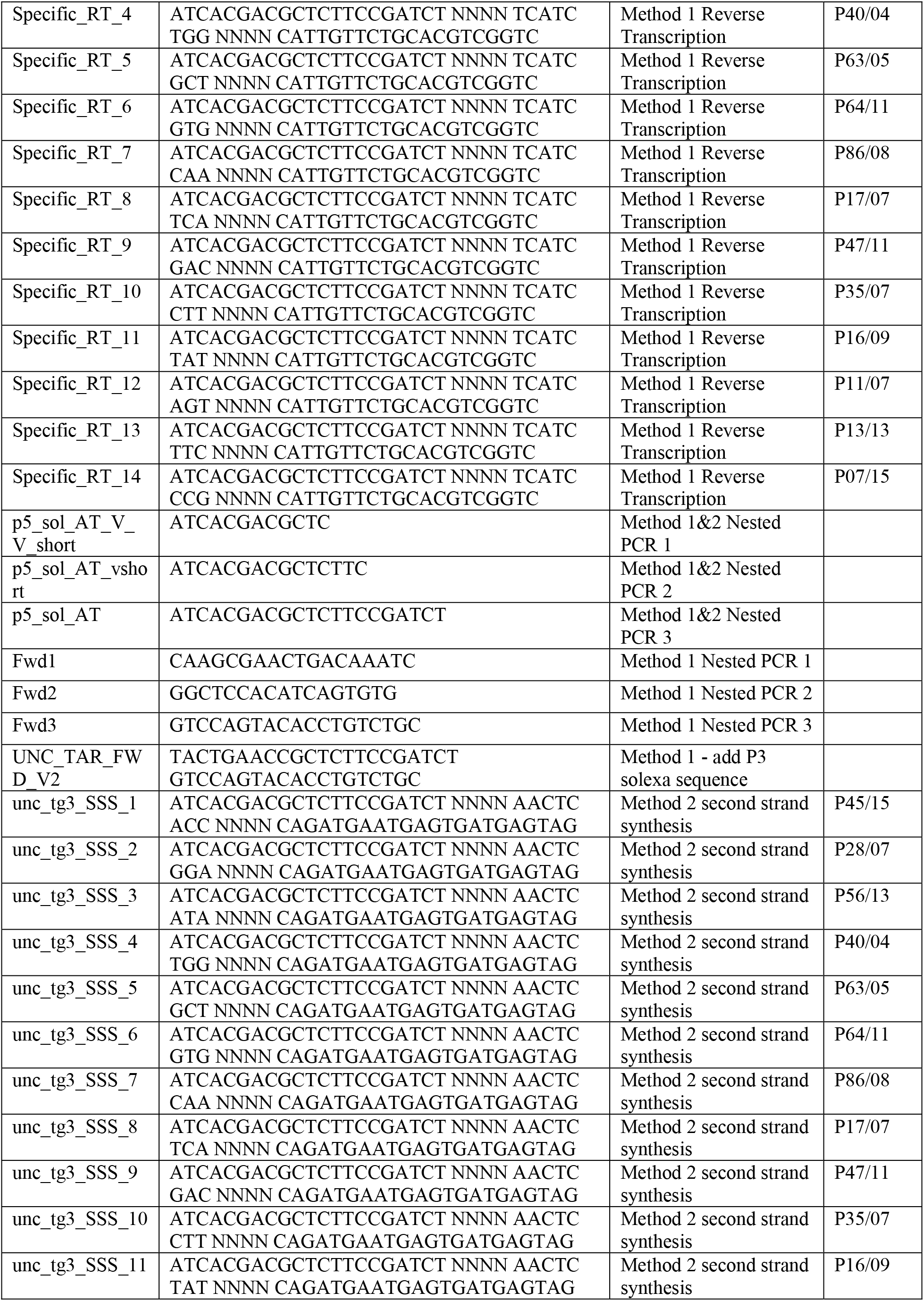
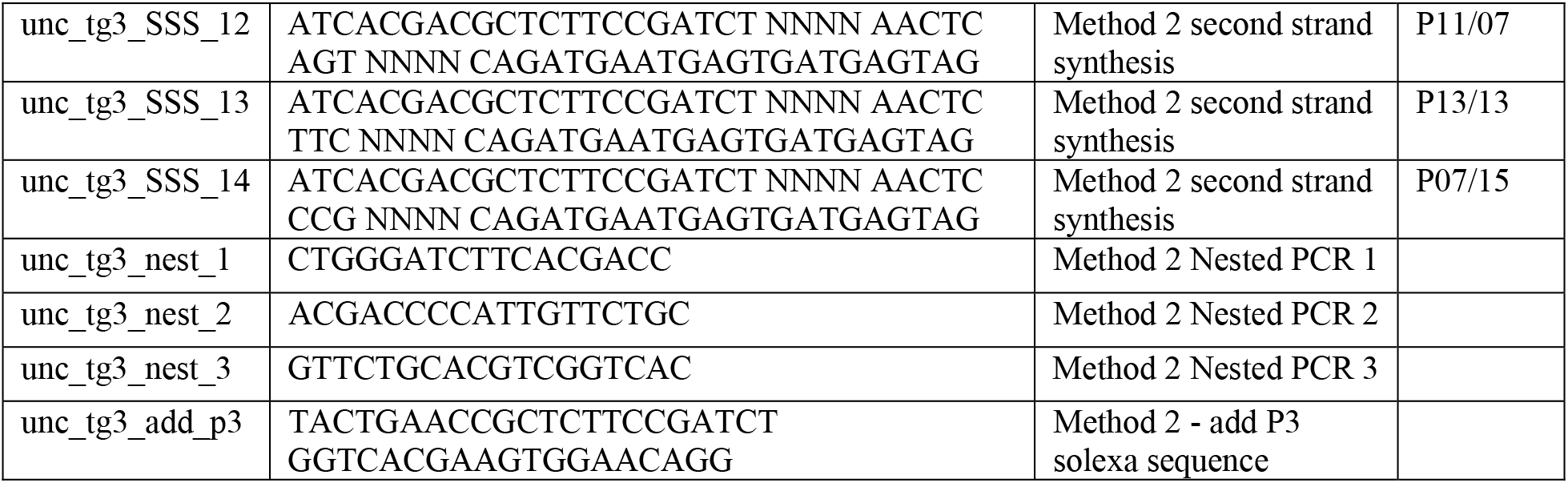

### Splicing reporters

One variant of the UNC13A exon 20, intron 20 and exon 21 sequence was synthesised and cloned into a pIRES-EGFP vector (Clontech) by BioCat. Plasmids with all four possible combinations of SNPs were generated by whole-plasmid PCR using primers with 5’ mismatches, followed by phosphorylation and ligation. Stbl3 bacteria grown at 30°C were used due to the observed instability of the plasmids in DH5alpha cells grown at 37°C. Sequences were verified by Sanger sequencing.

TDP-43 inducible knockdown SH-SY5Y cells were electroporated with 2 μg of DNA with the Ingenio electroporation kit (Mirus) using the A-023 setting on an Amaxa II nucleofector (Lonza). The cells were then left untreated or treated for 6 days with 1 μg/mL doxycycline before RNA extraction. Reverse transcription was performed with RervertAid (Thermo Scientific) and cDNA was amplified by nested PCR with miniGene specific primers 5’-TCCTCACTCTCTGACGAGG-3’ and 5’-CATGGCGGTCGACCTAG-3’ followed by UNC13A specific primers 5’-CAAGCGAACTGACAAATCTGCCGTGTCG-3’ and 5’- CGACACGGCAGATTTGTCAGTTCGCTTG-3’. PCR products were resolved on a TapeStation 4200 (Agilent) and bands were quantified with TapeStation Systems Software v3.2 (Agilent).

### TDP-43 protein purification

His-tagged TDP-43 was expressed in BL21-DE3 Gold *E. coli* (Agilent) as previously described (*57*). Bacteria were lysed by two hours of gentle shaking in lysis buffer (50 mM sodium phosphate pH 8, 300 mM NaCl, 30 mM imidazole, 1 M urea, 1% v/v Triton X-100, 5 mM beta-mercaptoethanol, with Roche EDTA-free cOmplete protease inhibitor) at room temperature. Samples were centrifuged at 16,000 rpm in a Beckman 25.50 rotor at 4°C for 10 minutes, and the supernatant was clarified by vacuum filtration (0.22 μm).

The clarified lysate was loaded onto a 5 ml His-Trap HP column (Cytiva) equilibrated with Buffer A (50 mM sodium phosphate pH 8, 300 mM NaCl, 20 mM imidazole) using an AKTA Pure system, and eluted with a linear gradient of 0-100% Buffer B (50 mM sodium phosphate pH 8, 300 mM NaCl, 500 mM imidazole) over 90 column volumes. The relevant fractions were then analysed by SDS-PAGE and then extensively dialysed (3.5 kDa cutoff) against ITC buffer (50 mM sodium phosphate pH 7.4, 100 mM NaCl, 1 mM TCEP) at 4°C.

### Isothermal titration calorimetry

RNAs with sequences 5’-AAGGAUGGAUGGAG-3’ (healthy) and 5’-AAGCAUGGAUGGAG-3’ (risk) were synthesised by Merck, resuspended in Ultrapure water, then dialysed against the same stock of ITC buffer overnight at 4°C using 1 kDa Pur-a-lyzer tubes (Merck). Protein and RNA concentrations after dialysis were calculated by A280 and A260 absorbance respectively. ITC measurements were performed on a MicroCal PEAQ-ITC calorimeter (Malvern Panalytical). Titrations were performed at 25°C with TDP-43 (9.6-12 μM) in the cell and RNA (96-120 μM) in the syringe. Data were analysed using the MicroCal PEAQ-ITC analysis software using nonlinear regression with the One set of sites model. For each experiment, the heat associated with ligand dilution was measured and subtracted from the raw data.

### iCLIP of minigene-transfected cells

HEK293T cells were transfected with either the 2x Healthy or 2x Risk minigenes using Lipofectamine 3000 (Thermofisher Scientific). Each replicate consisted of 2x 3.5 cm dishes, with two replicates per sample, for eight dishes total. 48 h after transfection, cells were crosslinked with 150 mJ/cm^2^ at 254 nm on ice, pelleted and flash frozen. Immunoprecipitations were performed with 4ug of TDP-43 antibody (proteintech 10782-2-AP) with 100ul of protein G dynabeads per sample, and iCLIP sequencing libraries were prepared as described in (*43*). Libraries were sequenced on an Illumina HiSeq4000 machine (SR100).

After demultiplexing the reads with Ultraplex, we initially aligned to the human genome using STAR (*32*), which showed that >5% of uniquely aligned reads mapped solely to the genomic region that is contained in the minigene. Given the high prior probability of reads mapping to the minigene, we therefore instead used Bowtie2 to align to the respective minigene sequences alone, thus minimising mis-mapping biases that could be caused by the SNPs(*44*) with settings “--norc --no-unal --rdg 50,50 --rfg 50,50 --score-min L,-2,-0.2 --end-to-end −N 1”, then filtered for reads with no alignment gaps, and length >25 nt. Due to the exceptional read depth and high library complexity, we did not perform PCR deduplication to avoid UMI saturation at signal peaks. All downstream analysis was performed using custom R scripts. Raw data is available at E-MTAB-10297.

## Supplementary Figures

**Fig. S1.**
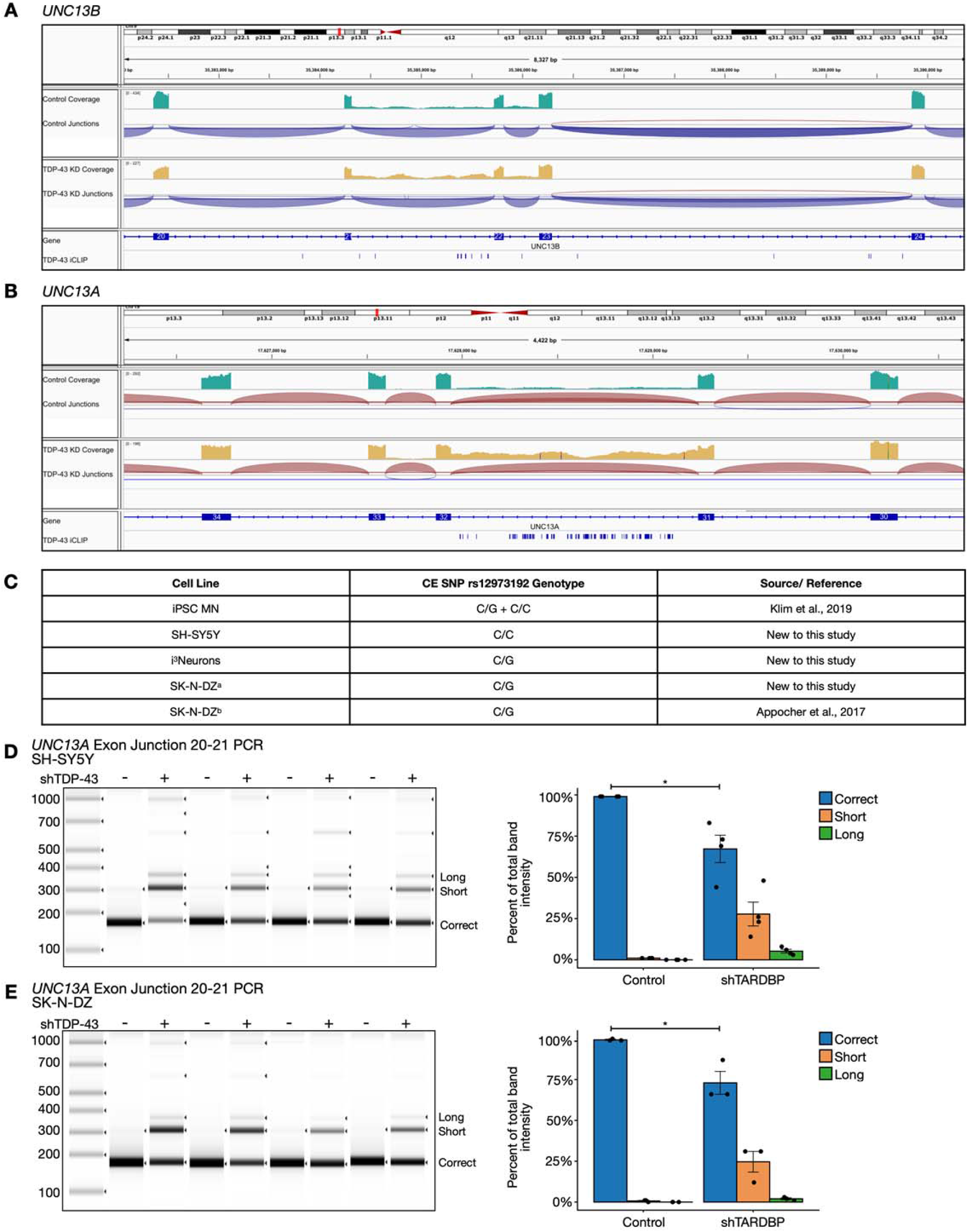
Characterization and validation of *UNC13A* and *UNC13B* mis-splicing after TDP-43 KD. **(A)** Genotypes and references of cell lines used in RNA-Seq analysis. **(B,C)** RNA-seq traces from IGV of representative samples from control (top) and *TARDBP* KD (bottom) in i^3^Neurons showing intron retention in *UNC13A* (A) (top) and *UNC13B* (B), overlaid with TDP-43 iCLIP peaks. **(D,E)** Capillary electrophoresis image of RT-PCR products for the *UNC13A* CE in SH-SY5Y (D) and SK-N-DZ (E) human cell lines upon *TARDBP* shRNA knockdown shows three products corresponding to correct splicing, a shorter and a longer CE. Replicates from each cell line are from 4 independent experiments. Graphs represent the means ± S.E. of the quantification of each PCR product as a percentage of the total product. N=4, Student’s t-test, *(p<0.05)

**Fig. S2.**
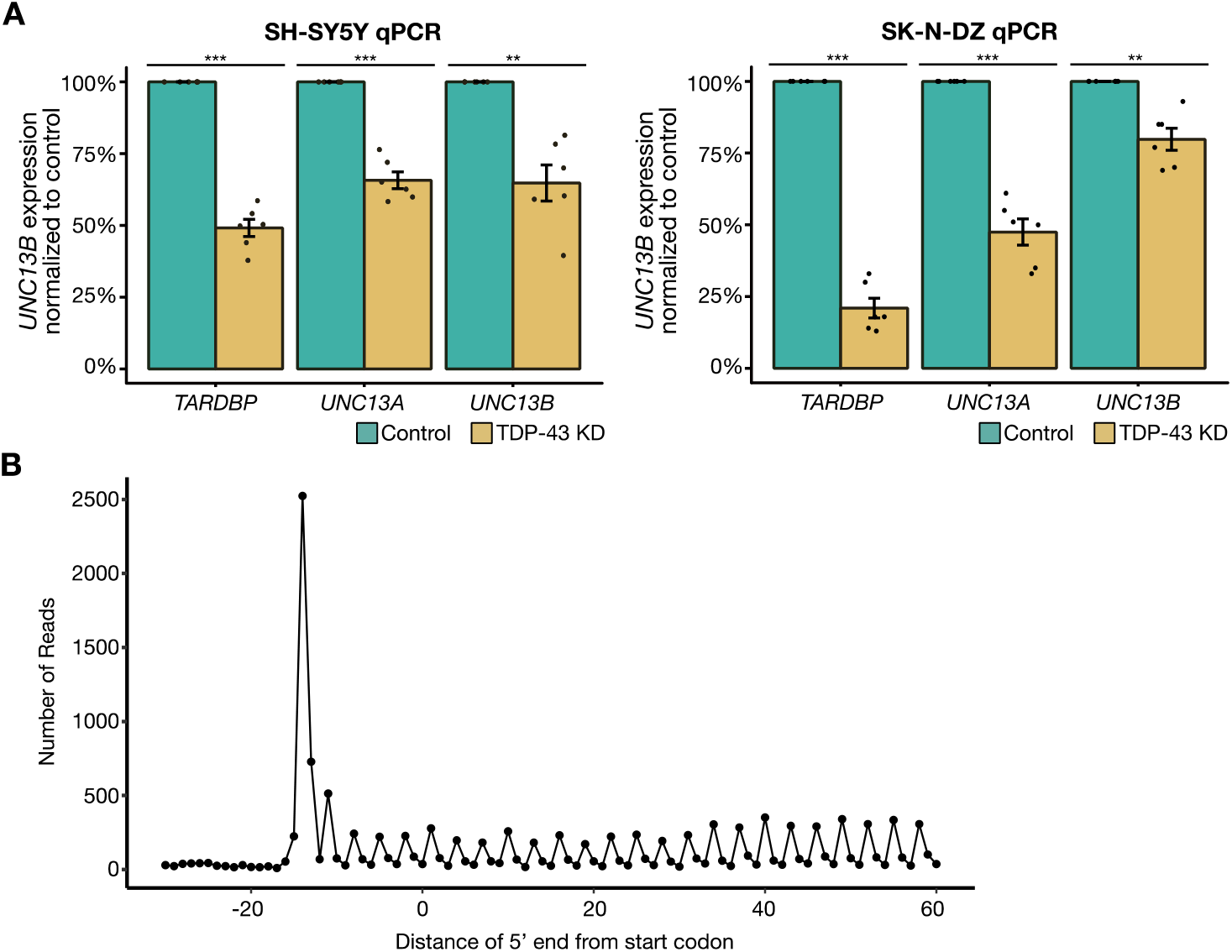
Orthogonal validation of *UNC13A* and *UNC13B* loss after TDP-43 KD. **(A)** RT-qPCR analysis shows *TDP-43, UNC13A* and *UNC13B* gene expression is reduced by *TARDBP* shRNA knockdown in both SH-SY5Y and SK-N-DZ human cell lines. Graphs represent the means ± S.E., N=6, One sample t-test, ***(p<0.001), **(p<0.01). **(B)** The 5’ ends of 29 nt reads relative to the annotated start codon from a representative ribosome profiling dataset (TDP-43 KD replicate B). As expected, we detected strong three-nucleotide periodicity, and a strong enrichment of reads across the annotated coding sequence relative to the upstream untranslated region.

**Fig. S3.**
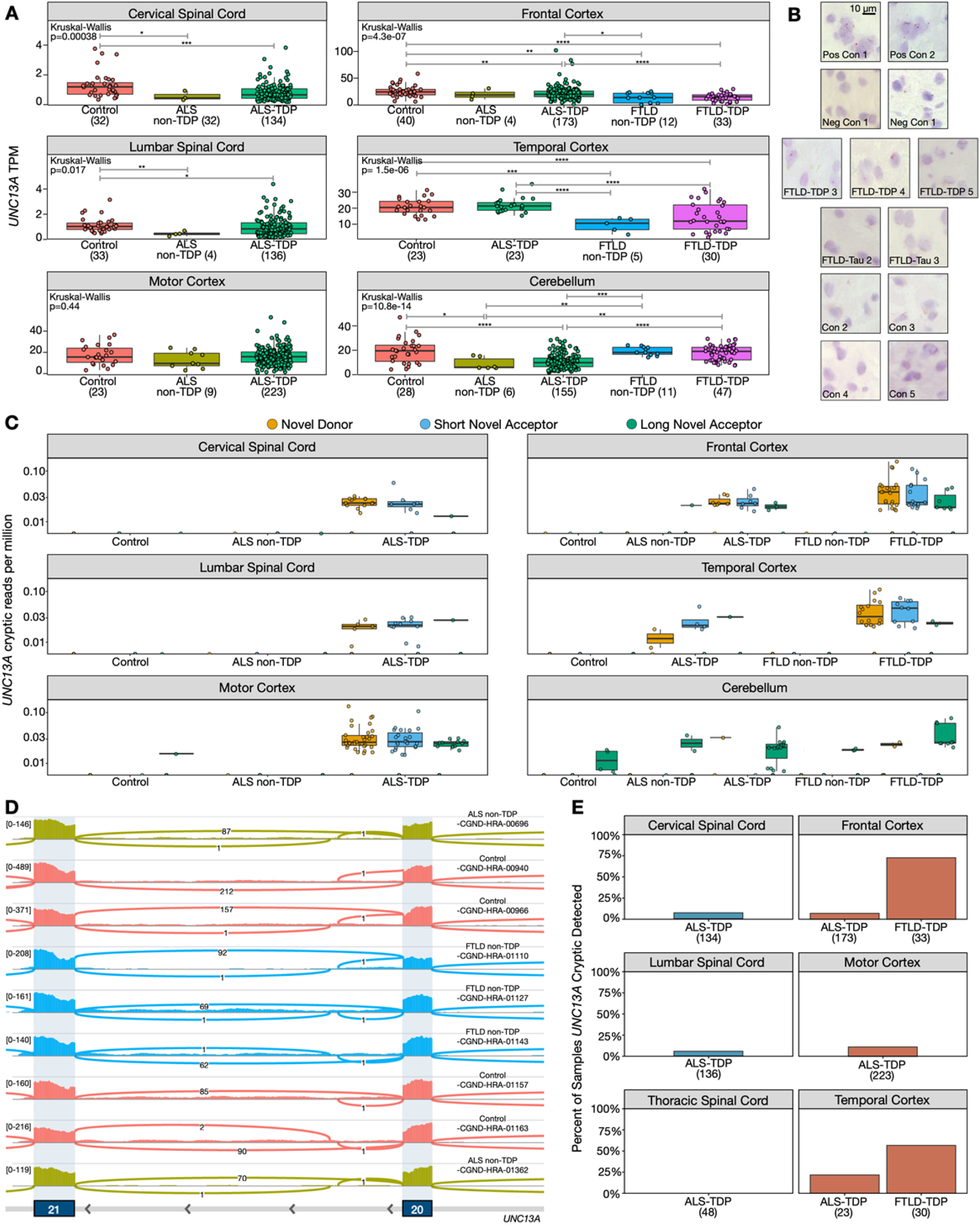
*UNC13A* transcript and CE expression across neuronal tissues. **(A)***UNC13A* expression across tissues and disease subtypes in the NYGC ALS Consortium RNA-seq dataset. Expression normalised as transcripts per million (TPM). Cortical regions have noticeably higher *UNC13A* expression than the spinal cord. **(B)** Representative images from RNA-based in situ hybridisation on frontal cortex brain tissue. Top and second rows show positive (PPIB-targeting) and negative (DapB-targeting) probe signal respectively. Other images are representative FTLD-TDP, FTLD-Tau and control cases validated with an UNC13a CE-targeting probe that were not shown in Fig. 3 **(C)** Expression of splice junction reads supporting the UNC13A CE across tissues and disease subtypes. Junction counts are normalised by library size in millions (junctions per million). The long novel acceptor junction is expressed across all disease subtypes in the cerebellum. **(D)** Example RNA-seq traces from IGV showing *UNC13A* cerebellar exon which shares the long novel acceptor junction as the *UNC13A* CE **(E)** Percentage disease relevant tissue samples with detectable *UNC13A* CE (1 supporting spliced read), split by disease and tissue.

**Fig. S4.**
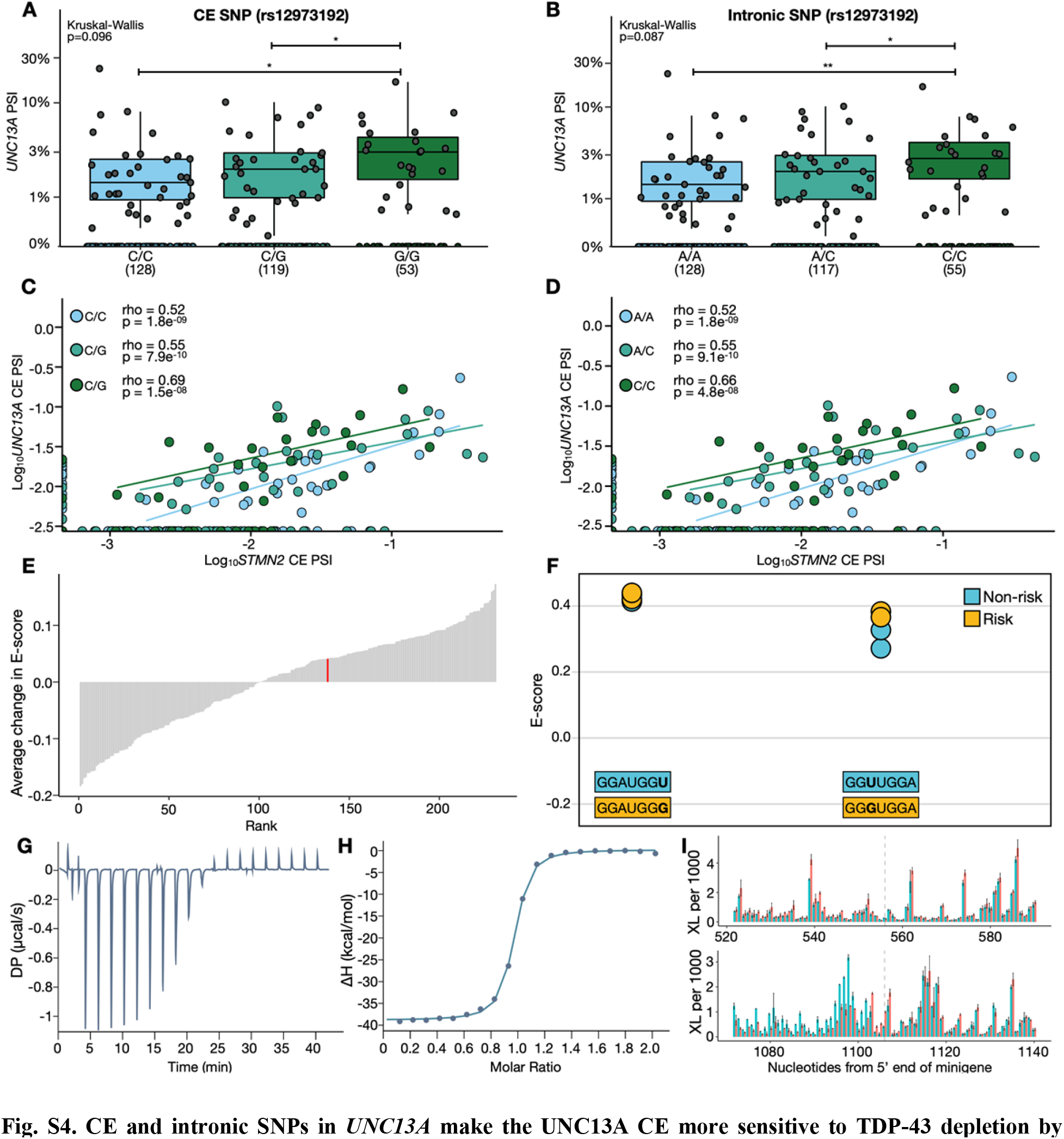
CE and intronic SNPs in *UNC13A* make the UNC13A CE more sensitive to TDP-43 depletion by altering TDP-43 binding affinity across the UNC13A CE-containing intron. **(A-B)***UNC13A* CE PSI by genotype (Wilcoxon test) **(C-D)** Effect of CE or intronic SNP on the correlation between *STMN2* and *UNC13A* CE PSI in ALS/FTD cortex in samples with at least 30 junction reads across the CE locus. **(E)** The average changes in E-score for heptamers containing the intronic SNP; red = TDP-43. **(F)** Individual TDP-43 E-scores for the two heptamers for which there was data (average change shown in red in (**D**)). Significance levels reported as *** (P<0.001); ** (P<0.01); * (P<0.05); ns (P > 0.05). **(G-H)** ITC measurement of the interaction of TDP-43 with 14-nt RNA containing the healthy sequence. A representative data set is reported, with raw data **(G)** and integrated heat plot **(H).** Circles indicate the integrated heat, the curve represents the best fit. **(I)** Mean crosslink density around the exonic (top) and intronic (bottom) SNPs in the 2H (red) and 2R (blue) minigenes, relative to the 5’ end of minigene (error bars = standard deviation; dashed lines show SNP positions).

**Table S1.**
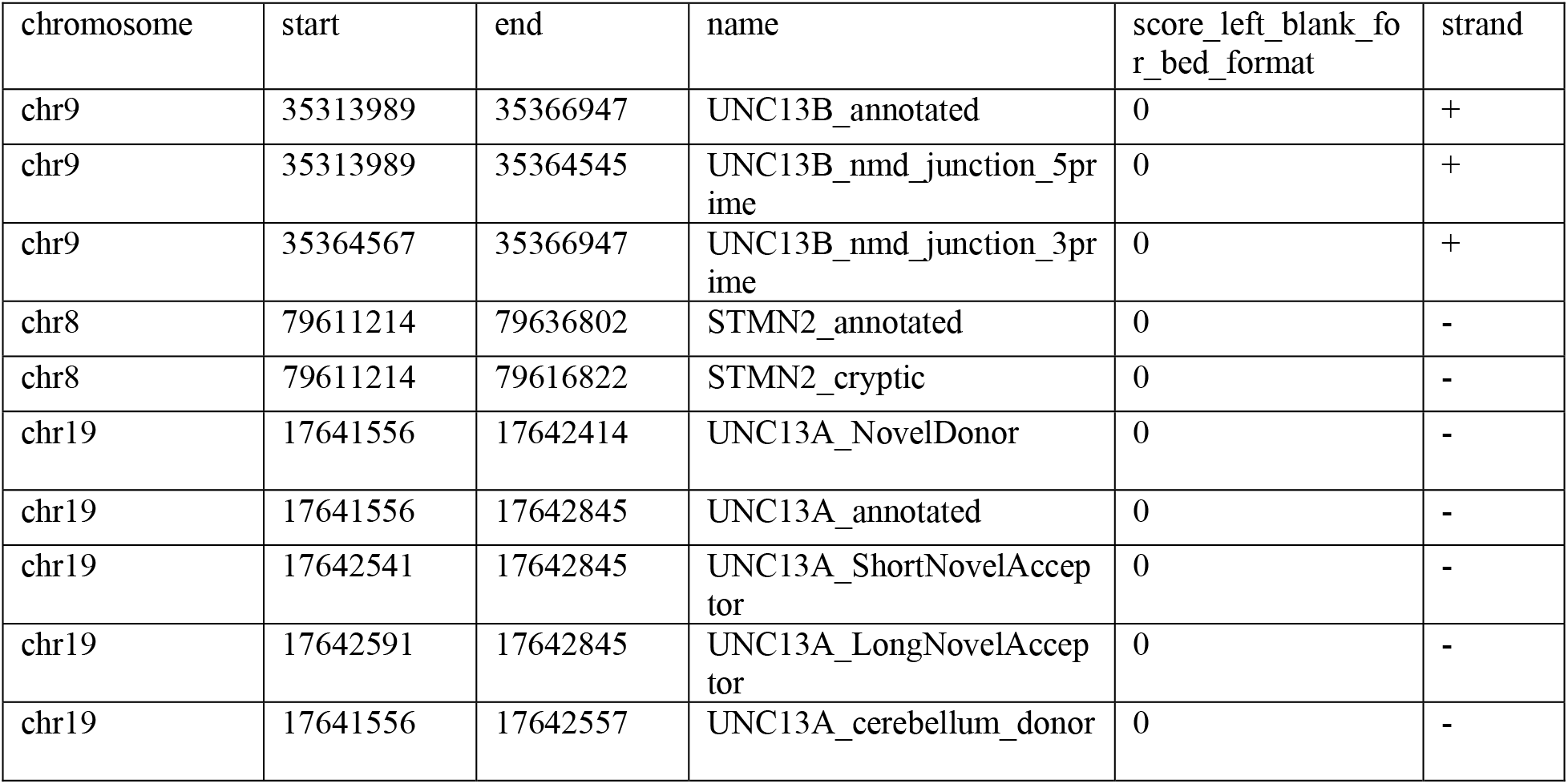
Hg38 coordinates for splice junctions used to calculate PSI.

### Data S1. (separate file)

List of differentially spliced junctions between control and TDP-43 KD i3Neurons (Fig. 1A).

### Data S2. (separate file)

List of differentially expressed genes between control and TDP-43 KD i3Neurons (Fig. 1B).

### Data S3. (separate file)

List of differentially ribosomal profiling genes between control and TDP-43 KD i3Neurons (Fig 2C).

### Data S4. (separate file)

Individual and average thermodynamic parameters obtained from ITC experiments

